# KOMPUTE: Imputing summary statistics of missing phenotypes in high-throughput model organism data

**DOI:** 10.1101/2023.01.12.523855

**Authors:** Coby Warkentin, Michael J. O’Connell, Donghyung Lee

## Abstract

**Motivation:** The International Mouse Phenotyping Consortium (IMPC) is striving to build a comprehensive functional catalog of mammalian protein-coding genes by systematically producing and phenotyping gene-knockout mice for almost every protein-coding gene in the mouse genome and by testing associations between gene loss-of-function and phenotype. To date, the IMPC has identified over 90,000 gene-phenotype associations, but many phenotypes have not yet been measured for each gene, resulting in largely incomplete data; about 75.6% of association summary statistics are still missing in the latest IMPC summary statistics dataset (IMPC release version 16).

**Results:** To overcome these challenges, we propose KOMPUTE, a novel method for imputing missing summary statistics in the IMPC dataset. Using conditional distribution properties of multivariate normal, KOMPUTE estimates association Z-scores of unmeasured phenotypes for a particular gene as a conditional expectation given the Z-scores of measured phenotypes. We evaluate the efficacy of the proposed method for recovering missing Z-scores using simulated and real-world data sets and compare it to a singular value decomposition (SVD) matrix completion method. Our results show that KOMPUTE outperforms the comparison method across different scenarios.

**Availability and implementation:** An R package for KOMPUTE is publicly available at https://github.com/statsleelab/kompute, along with usage examples and results for different phenotype domains at https://statsleelab.github.io/komputeExamples.

**Supplementary information:** Supplementary data are available at *Bioinformatics* online.

## 1 Introduction

The International Mouse Phenotyping Consortium (IMPC) has been cataloging the functions of the entire mouse genome by producing knockout mouse lines for all protein-coding genes and examining the effects on various behavioral, physiological, morphological, and biochemical phenotypes. (Haselimashhadi, et al., 2020; Meehan, et al., 2017). This process to date has successfully identified over 90,000 gene-phenotype associations in a controlled and reproducible setting (IMPC release version 16). However, generating a comprehensive functional catalog of the mouse genome is a daunting task, as there are many potential gene-phenotype pairs to consider, and testing all of them in a controlled setting is time-consuming and resource-intensive. Furthermore, knocking out vital genes can often result in early embryonic lethality or developmental abnormalities, making many phenotypes unmeasurable for mice from which these genes are removed (Dickinson, et al., 2016). Therefore, many phenotypes of interest have not yet been measured for many genes, so the IMPC association summary statistics data is still largely incomplete (Figure 1).

**Figure 1.**
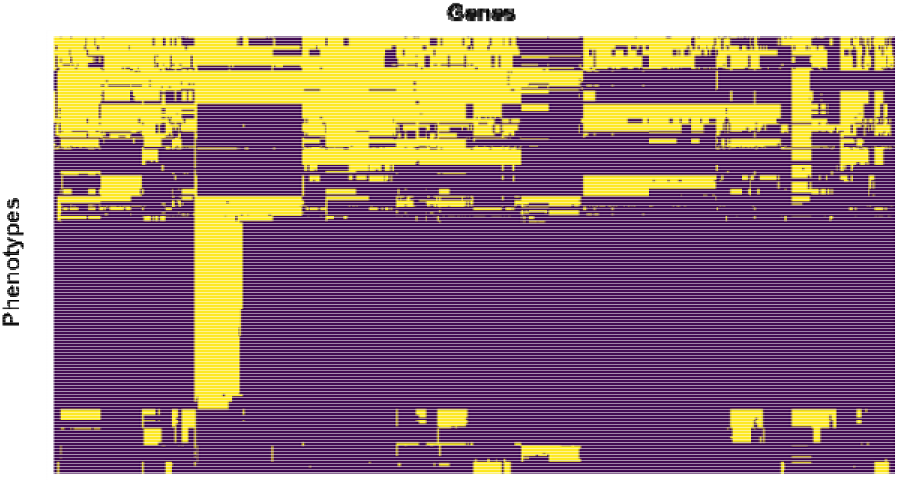
Heatmap showing the presence or absence of data for each gene-phenotype pair in the IMPC data (release version 16). Dark purple cells indicate missing data, whereas light yellow cells indicate measured data. Each column represents one of 8216 genes that have been knocked out in various mice, and each row represents one of 303 phenotypes. Overall, only about 24.4% of gene-phenotype pairs are tested.

To effectively recover summary statistics of missing phenotypes, we propose a novel method, KOMPUTE. KOMPUTE uses the conditional distribution properties of multivariate normal to directly impute association statistics (i.e., two-sided association Z-scores) of missing phenotypes in the IMPC data. A similar idea has been used to directly impute association Z-scores of unmeasured genetic variants in genome-wide association studies while still maintaining high imputation accuracy and reducing computational burden significantly (Lee, et al., 2013; Lee, et al., 2015). Similarly, the KOMPUTE method allows missing association Z-scores to be imputed directly, avoiding the intermediate step of having to impute or measure missing phenotypes for each mouse subject.

## 2 Methods

### 2.1 Summary statistics imputation

KOMPUTE estimates association Z-scores of unmeasured phenotypes using well-known conditional expectation formulas of the multi-variate normal distribution (Lee, et al., 2013). Assume that the summary statistics capturing the association between all genes and phenotypes form an matrix of two-sided association Z-scores, where is the total number of phenotypes observed and is the total number of genes tested. For a particular gene, let be the vector of the unmeasured Z-scores (length, i.e., number of missing phenotypes =) and let be the vector of the measured Z-scores (length). By reordering phenotypes in this way, each column vector of Z-scores can be written as

Let be the genetic correlation matrix between phenotypes, where the entry in the -th row and -th column represents the genetic correlation between phenotypes and according to the new ordering, . Then can be written as

where is the genetic correlation matrix of unmeasured pheno-types, and are the and genetic correlation matrices between unmeasured and measured phenotypes, respectively, and is the genetic correlation matrix of measured phenotypes.

Under the null hypothesis of no association between gene-knockout and phenotype, asymptotically follows the multivariate normal distribution, where is the genetic correlation matrix between phenotypes. Using this, can be estimated as

The corresponding variance-covariance matrix is then

The diagonal of this matrix represents the uncertainty of estimator. Therefore, the diagonal elements of can be used as imputation

accuracy measure (imputation information) of. The imputation information value ranges from 0 to 1 for the corresponding imputed Z-score, with values closer to 1 indicating less variation in the imputed estimate and therefore a more reliable estimate. To ensure that is invertible, a small ridge penalty is added to each diagonal element of. The ridge penalty added ( ) is chosen to be small enough to have minimal impact on the imputed Z-scores (Lee, et al., 2015).

By repeating this method for each gene in the original association summary statistics matrix, we can generate a new matrix with imputed Z-scores for the previously unmeasured phenotypes.

### 2.2 Estimating phenotypic correlation as a proxy for genetic correlation

Genetic correlation refers to the amount of variance shared between two phenotypes due to genetic factors. Estimating genetic correlations accurately can be difficult because it often requires collecting genetic information from very large samples, particularly when the heritability of the two phenotypes is low (Sodini, et al., 2018). However, previous studies (Reusch and Blanckenhorn, 1998; Roff, 1995; Sodini, et al., 2018; Waitt and Levin, 1998) have found empirical evidence of a strong similarity between genetic and phenotypic correlations in insets, plants, animals, and humans when the sample size is large, based on the conjecture initially proposed by James Cheverud in 1988 (Cheverud, 1988). Based on this evidence, we use the easier-to-estimate phenotypic correlation matrix observed in control mice as a proxy for the genetic correlation matrix ( ) in the imputation method.

The raw phenotype data collected in experimental settings often affected by non-biological factors (e.g., phenotyping center) that can significantly impact measured phenotypes. In order to correct for these potential confounding factors, we use principal variance component analysis (PVCA) (Chen, et al., 2011; Li, et al., 2009) to identify the proportion of phenotypic variation explained by non-biological covariates and Combat (Johnson, et al., 2007) to remove the effects of the covariates from the phenotype data. The resulting adjusted phenotype data is then used to calculate the phenotypic correlation matrix using the Pearson correlation coefficient.

### 2.3 Simulation studies

To evaluate the performance of our proposed imputation method, KOMPUTE, we conducted extensive simulation studies and compared its performance to that of a singular value decomposition (SVD) matrix completion method (Kurucz, et al., 2007). We simulated 100 association Z-score matrices (10,000 genes by 8 phenotypes) from a multivariate normal distribution), where denotes the phenotype correlation matrix estimated from the eight body composition phenotypes of control mice.

We compared three different imputation methods: SVD matrix completion, KOMPUTE considering all imputed values, and KOMPUTE while only considering values with sufficiently high imputation information (above 0.8). Each of these methods was tested under three different scenarios where 20%, 40%, and 60% of the simulated Z-scores were randomly masked (i.e., temporarily deleting them from the data) and imputed. The Pearson correlation coefficient between masked Z-scores and imputed Z-scores was computed for each scenario and averaged across the 100 simulations.

In order to assess the performance of the proposed imputation method in realistic scenarios, we conducted additional experiments using real measured association Z-scores of three phenotype domains: body composition (8 phenotypes), clinical chemistry (19 phenotypes), and open field (14 phenotypes). For each domain, we first estimated a phenotype correlation matrix using control mice phenotypes, adjusting for non-biological factors through PVCA and Combat.

We then randomly masked 1000 gene-phenotype association Z-scores from each of the three domains and used the remaining Z-scores and the estimated phenotype correlation matrix to impute the missing values with the KOMPUTE method. We considered only the imputed Z-scores with high imputation information (greater than 0.8) and calculated the Pearson correlation coefficient between imputed and original Z-scores as a measure of the effectiveness of KOMPUTE.

## 3 Results

The KOMPUTE method demonstrated superior performance compared to the SVD method across all simulation scenarios (Table 1). Furthermore, restricting our analysis to imputed Z-scores with high imputation information (i.e., KOMPUTE w/ info > 0.8) resulted in even more reliable results. These highly accurate imputed values were obtained even when a significant proportion of the simulated data was missing.

**Table 1.**
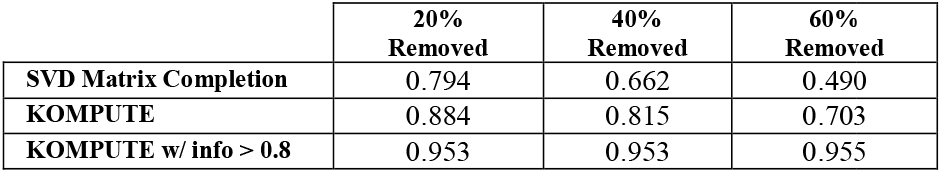
Pearson correlation coefficients between the original and imputed Z-scores. Three imputation methods were each tested under three different scenarios in which 20%, 40%, or 60% of the Z-scores were removed and imputed. The mean correlations over 100 simulations are presented, with a standard deviation less than 0.01 for all estimates.

Figure 2 shows the KOMPUTE method’s effectiveness in imputing missing Z-scores in realistic scenarios. In the body composition domain, all 1000 imputed Z-scores were complete cases with valid imputation information. The Pearson correlation between the original Z-scores and the imputed Z-scores was 0.793. Setting a cutoff for the imputation information at 0.8 to only include estimates with a higher degree of confidence resulted in 583 of the 1000 Z-scores being kept, with a Pearson correlation of 0.841 between these scores and the original masked Z-scores. For the clinical chemistry domain, all 1000 imputed Z-scores were complete and valid, but many of the imputation information values were relatively low, leading to a Pearson correlation of 0.486 between the masked and imputed Z-scores. However, the 113 cases where the information was greater than 0.8 had a Pearson correlation of 0.874 between the masked and imputed Z-scores. Finally, for the open field domain, all 1000 Z-scores were valid and generally had high imputation information values, resulting in a Pearson correlation of 0.929 between masked and imputed Z-scores. When the threshold for imputation information was set at 0.8, 846 out of the 1000 Z-scores met this criterion, and these imputed Z-scores showed a very strong Pearson correlation of 0.961 with the original masked Z-scores.

**Figure 2.**
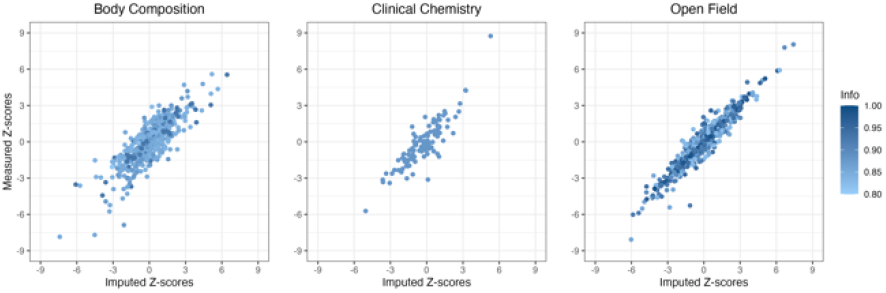
Measured Z-scores as a function of imputed Z-scores for three phenotype domains, namely body composition, clinical chemistry, and open field. A total of 1,000 Z-scores were masked in each domain and imputed using the KOMPUTE method. To ensure the reliability of the imputed values, an imputation information score of at least 0.8 was required. The recovery rates for the body composition, clinical chemistry, and open field domains were 58.3%, 11.3%, and 84.6%, respectively, based on this threshold. The Pearson correlation coefficients between the original and imputed Z-scores for these domains were 0.841, 0.874, and 0.961, respectively.

## 4 Discussion and conclusion

KOMPUTE has shown to be an effective method for imputing missing summary statistics in high-throughput model organism-based genetic association studies such as IMPC studies. The method outperformed the SVD matrix completion in all simulation scenarios and demonstrated good performance in more realistic settings. Furthermore, the use of an imputation information score allows for the identification of more reliable estimates. By improving the completeness of the IMPC dataset, KOMPUTE can help researchers unlock the full potential of this valuable resource. KOMPUTE also has the potential to be applied to a variety of high-throughput studies in various other model organisms, including Caenorhabditis elegans or Arabidopsis thaliana, where comprehensive phenotyping is a significant challenge.

## Acknowledgements

The authors thank the International Mouse Phenotyping Consortium for sharing summary statistics and mouse phenotype data sets.

